# Automatic encoding of a view-centered background image in the macaque temporal lobe

**DOI:** 10.1101/2020.03.01.971507

**Authors:** He Chen, Yuji Naya

## Abstract

Perceptual processing along the ventral visual pathway to the hippocampus is hypothesized to be substantiated by signal transformation from retinotopic space to relational space, which represents interrelations among constituent visual elements. However, our visual perception necessarily reflects the first person’s perspective based on the retinotopic space. To investigate this two-facedness of visual perception, we compared neural activities in the temporal lobe (anterior inferotemporal cortex, perirhinal and parahippocampal cortices, and hippocampus) between when monkeys gazed on an object and when they fixated on the screen center with an object in their peripheral vision. We found that in addition to the spatially invariant object signal, the temporal lobe areas automatically represent a large-scale background image, which specify the subject’s viewing location. These results suggest that a combination of two distinct visual signals on relational space and retinotopic space may provide the first person’s perspective serving for perception and presumably subsequent episodic memory.

## Introduction

Visual information of our external world could be once decomposed into “what” and “where” before we attained its mental representation as the first person’s perspective (Eichenbaum, Yonelinas, & Ranganath, 2007; Palombo et al., 2015; Tulving, 2002). For several decades, it has been considered that the perception of these two visual features proceeds exclusively through the ventral and dorsal pathways (Goodale & Milner, 1992; Haxby et al., 1991; Mishkin & Ungerleider, 1982). Instead of this widespread dichotomy, contemporary visual neuroscience research suggests a presence of spatial information in the ventral pathway for perception (Chen & Naya, 2019; Connor & Knierim, 2017; Russell A. Epstein & Julian, 2013; Freud, Plaut, & Behrmann, 2016; Hong, Yamins, Majaj, & DiCarlo, 2016; Kornblith, Cheng, Ohayon, & Tsao, 2013; Mormann et al., 2017; Schenk, 2010). For instance, neurons in the inferotemporal (IT) cortex (TEO and TEd) of non-human primates exhibited preferential responses to scene-like stimuli rather than object-like stimuli (Kornblith et al., 2013; Vaziri, Carlson, Wang, & Connor, 2014). The response pattern of scene-selective IT neurons may be comparable to an activation pattern in the parahippocampal place area detected in human functional imaging studies (R. Epstein & Kanwisher, 1998; Julian & Epstein, 2013). The parahippocampal place area is located within the parahippocampal cortex (PHC) of the medial temporal lobe (MTL), which receives inputs from the early stages of the ventral pathway including the TEO and posterior TEd in addition to inputs from the dorsal pathway, and provides spatial information to the hippocampus (HPC) – a candidate of the final brain region for scene perception (Burgess, 2008) - via the medial/posterior entorhinal cortex (ERC) (Rolls, 2018). On the other hand, neurons in the IT cortex also represent location information of an object within a scene either at population-coding level (Hong et al., 2016) or at single-neuron level (Chen & Naya, 2019). It is worth noting that while most neurophysiological studies had shown a spatial invariance of object at single-neuron level during monkeys’ fixating on the center of a display under either the passive-viewing task (Hong et al., 2016; Kobatake & Tanaka, 1994) or delayed matching-to-sample (object) task (Miyashita & Chang, 1988; Nakamura, Matsumoto, Mikami, & Kubota, 1994), our recent study demonstrated equivalent or even more neurons exhibiting location signal compared with object signal in the ventral part of the anterior inferotemporal cortex (TEv) and its downstream MTL area (e.g., perirhinal cortex, PRC) during an item-location retention (ILR) task requiring monkeys to encode both identity and location of a sample object using a foveal vision (Fig. 1). Importantly, the location-selective activity during the ILR task could not be explained by the animals’ eye-positions themselves (Chen & Naya, 2019).

**Fig. 1.**
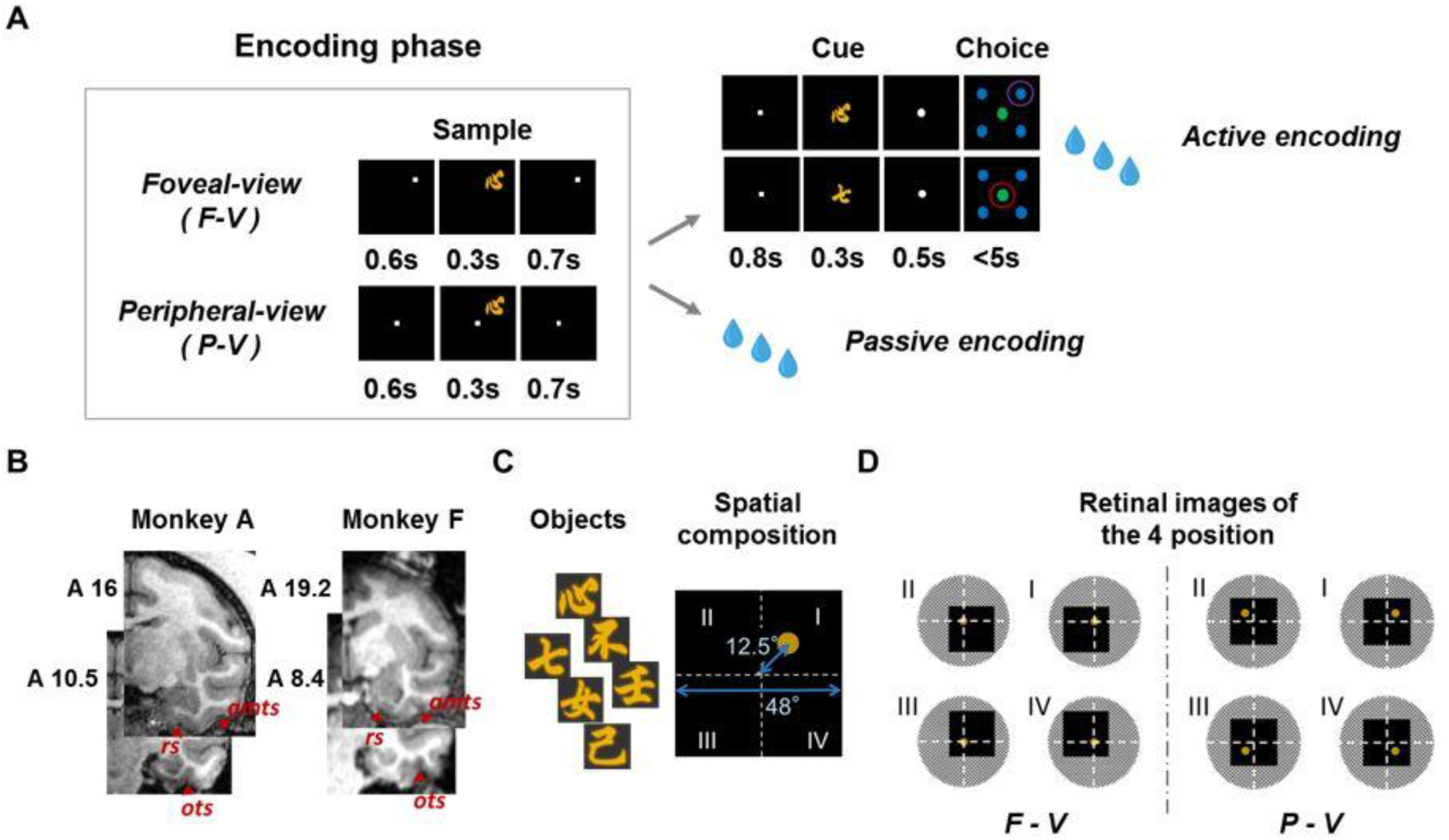
Encoding of location and item in two view conditions. (A) Schematic diagram of location and item encoding in the F-V and P-V conditions of the active-encoding and passive-encoding tasks. In the active-encoding task, the cue stimulus was the same as the sample stimulus during the encoding phase in the match trial (Top), while the two stimuli differed in the nonmatch trial (Bottom). Red circles indicate correct answers. Passive-encoding task consisted of only the encoding phase of the active-encoding task. (B) Example of coronal sections from monkey A and monkey F. The sections from Monkey A are 16 mm and 10.5 mm anterior to the interaural line and include the hippocampus (HPC), parahippocampal cortex (PHC), perirhinal cortex (PRC), and area TE (TE). amts, anterior middle temporal sulcus; ots, occipital temporal sulcus; rs, rhinal sulcus. Coronal sections from monkey F are 19.2 mm and 8.4 mm anterior to the interaural line. (C) Six object stimuli were used in the task, and an example of spatial composition during the sample period is shown. A yellow disk indicates an object position. (D) Schematic diagram of visual inputs to the retinae during the sample period; white dashed lines indicate the horizontal and vertical meridians of the visual field.

Considering that different gaze positions cause a substantial difference in the large-scale visual input in the ILR task using the foveal-view (F-V) condition (Fig. 1D), the most straightforward explanation for the robust location signal might be that a substantial number of neurons in the IT cortex and MTL areas are driven by the retinotopic signal including parafoveal vision, which would not only serve for recognizing a scene (Connor & Knierim, 2017; Dilks, Julian, Kubilius, Spelke, & Kanwisher, 2011; Kornblith et al., 2013; Vaziri et al., 2014) but also signal a particular location in the scene (Chen & Naya, 2019; Hong et al., 2016). An alternative explanation would be the location information of an object is coded into internal spatial relationships within a large complex stimulus including an object and its background regardless of their absolute retinotopic positions. In other words, the IT cortex and MTL areas would represent object location by transforming representations of the object and its background on the retinotopic space (Zhaoping, 2019) into those on the “relational space” (Connor & Knierim, 2017). In this case, the location signal in the ILR task would be sensitive to the task demand requiring the animals to retain the object location for a following action rather than a retinotopic image depending on the animals’ gaze position.

To address this question and investigate characteristics of spatial information in the ventral pathway and its downstream (i.e., MTL areas), we examined single-unit activities and local-field potentials (LFPs) from the TEv and MTL subregions during an object stimulus presented randomly at one of the quadrants on the display in a peripheral-view (P-V) as well as in an F-V condition (Fig. 1). In the P-V condition, animals were required to fixate on a central dot and obtain the location and item-identity information of the sample object using their peripheral vision (Fig. 1A). We compared the location effects between the two-view conditions by testing two rhesus macaques, and found that regardless of the task demands for encoding of an object and its location, there were much more abundant location signal in the F-V condition compared with the P-V condition on all the recording regions of the two monkeys.

## Results

We collected data in both F-V and P-V conditions from two rhesus macaques (Fig. 1B). During the recording, Monkey A was required to encode an identity of a sample stimulus and its location actively for a subsequent response (i.e., ILR task). We reported the single-unit data in the F-V condition of the ILR task in the previous study (Chen & Naya, 2019); here, we refer to the ILR task as an “active-encoding task.” On the other hand, monkey F was only required to fixate on a small white dot, viewing a sample stimulus passively (“passive-encoding task”) in both view conditions (Figs. 1A&B). We did not record from the single monkeys in both encoding tasks because it would be difficult to exclude both explicit and implicit influences of learning the active-encoding task on the cognitive process in the passive-encoding task. We used the same six visual objects (yellow Chinese characters, radius = 3°) as sample stimuli for both monkeys through all the recording sessions (Fig. 1C). It should be noted that the retinotopic images differed entirely between F-V and P-V conditions although a position of a sample stimulus was identical relative to the external world including a large square background (48° each side) on the display between the two view conditions (Figs. 1C&D). This two-by-two experimental design (“F-V vs. P-V” × “active-encoding vs. passive-encoding”) allowed us to compare the neural signals in the F-V condition with those in the P-V condition in the animals with different task demands. Monkey A performed the active-encoding task at high performances in both F-V (96.2 ± 3.7 %, 454 sessions) and P-V (92.2 ± 7.1 %, 477 sessions) conditions.

### Gaze-related location signal

We first investigated single-unit activities signaling location information. Figure 2A shows an example of TEv neurons that were recorded in the active-encoding task. The neuron showed the largest responses when the animal fixated on the *position I* (*top right* on the large square background). Although the responses once decayed, the neuron responded strongly when an item stimulus was presented as a sample stimulus at the same *position I* in the F-V condition. We examined the neuronal responses during 80-1000 ms after the onset of sample presentation (sample period) using a two-way ANOVA with item identities (six items) and locations (four locations) as main effects. The neuron showed a significant location effect [*P* < 0.0001, *F*(3,156) = 18.98] but not for item identities of sample stimuli [*P* = 0.309, *F*(5,156) = 1.21]. In contrast to the strong location-selectivity in the F-V condition, the same TEv neuron did not show location-selective activities in the P-V condition during the sample period [*P* = 0.183, *F*(3,157) = 1.63].

**Fig. 2.**
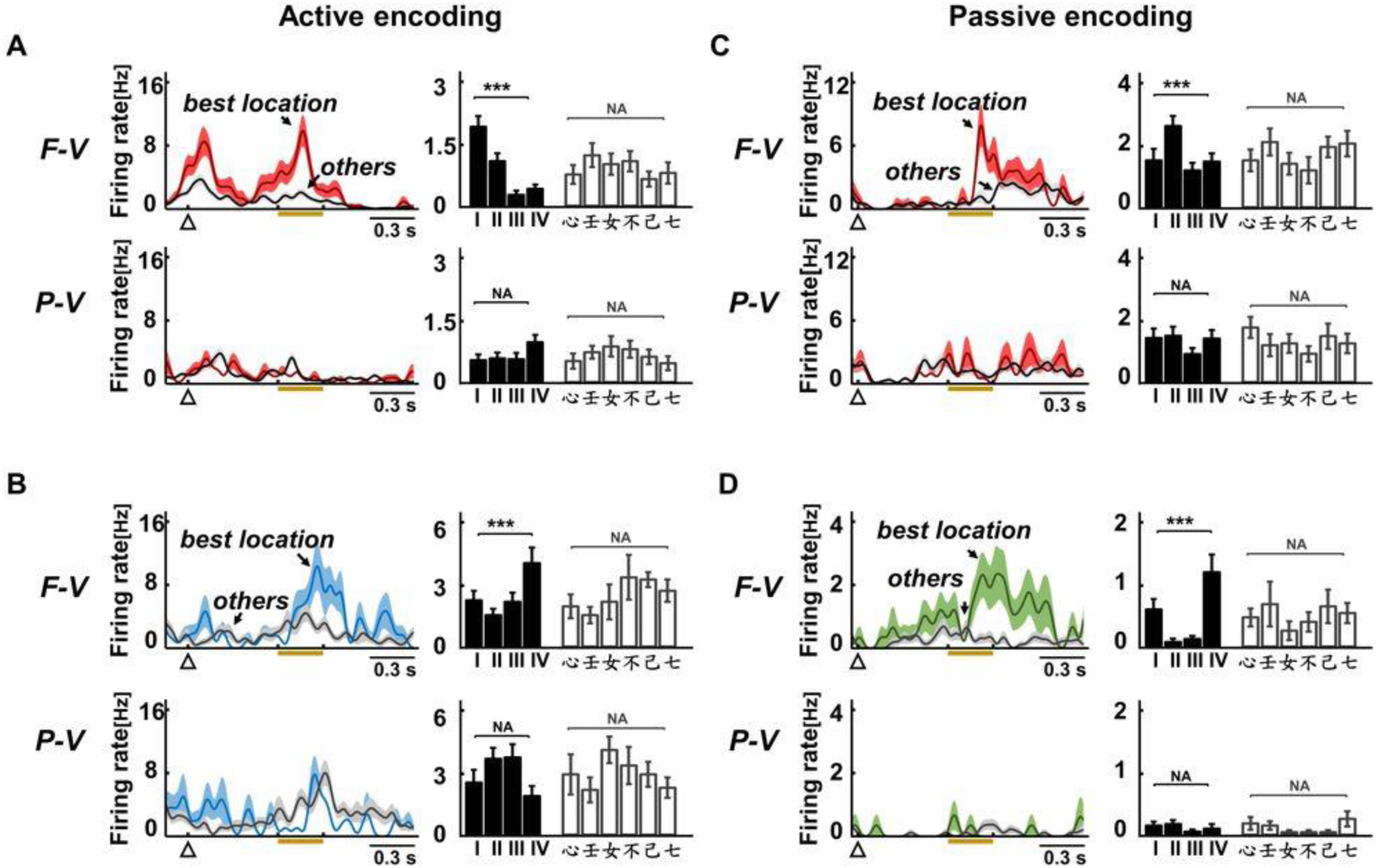
Responses of the location-selective cells in the active-encoding and passive-encoding task. (A) Example of the location-selective cells from TE in the F-V and P-V condition of the active-encoding task. (Left) Spike-density functions (SDFs) (sigma = 20 ms) indicating the firing rates under two conditions (best location and the average of other three locations). (Right) Bar graph indicating the mean firing rate during sample period (80-1000 ms after sample on) under each location and each item. (B) Example of the location-selective cells from PRC in the F-V and P-V condition of the active-encoding task. (C-D) Examples of the location-selective cells in TE (C) and PHC (D) in the F-V and P-V conditions of the passive-encoding task.

Figure 2B shows an example of PRC neurons that also exhibited location-selective activities only in the F-V conditions. This neuron signaled location information only after sample presentation in the F-V condition, suggesting that the presence of location signal in the F-V condition cannot be necessarily explained by preceding location-selective activity before sample presentation. We examined the prevalence of location signal in the two view conditions among the recording regions by calculating proportions of neurons with significant (*P* < 0.01, two-way ANOVA) location-selective activities during the sample period in each area. All recording regions contained significantly (*P* < 0.0016 in each region, *χ*^2^ test) larger proportions of location-selective cells in the F-V condition (24%, TE; 27%, PRC; 21%, HPC; 20%, PHC) than the P-V condition (7%, TE; 10%, PRC; 7%, HPC; 4%, PHC) (Fig. 3A). These results indicated that the location information in the ventral pathway and MTL areas were sensitive to the view conditions, although the same task-relevant information was required for a following action in the active-encoding task. The robust location signal only in the F-V condition implicates that the temporal lobe areas represent a visual image, which subjects view rather than the goal-directed spatial information related with an action plan.

**Fig. 3.**
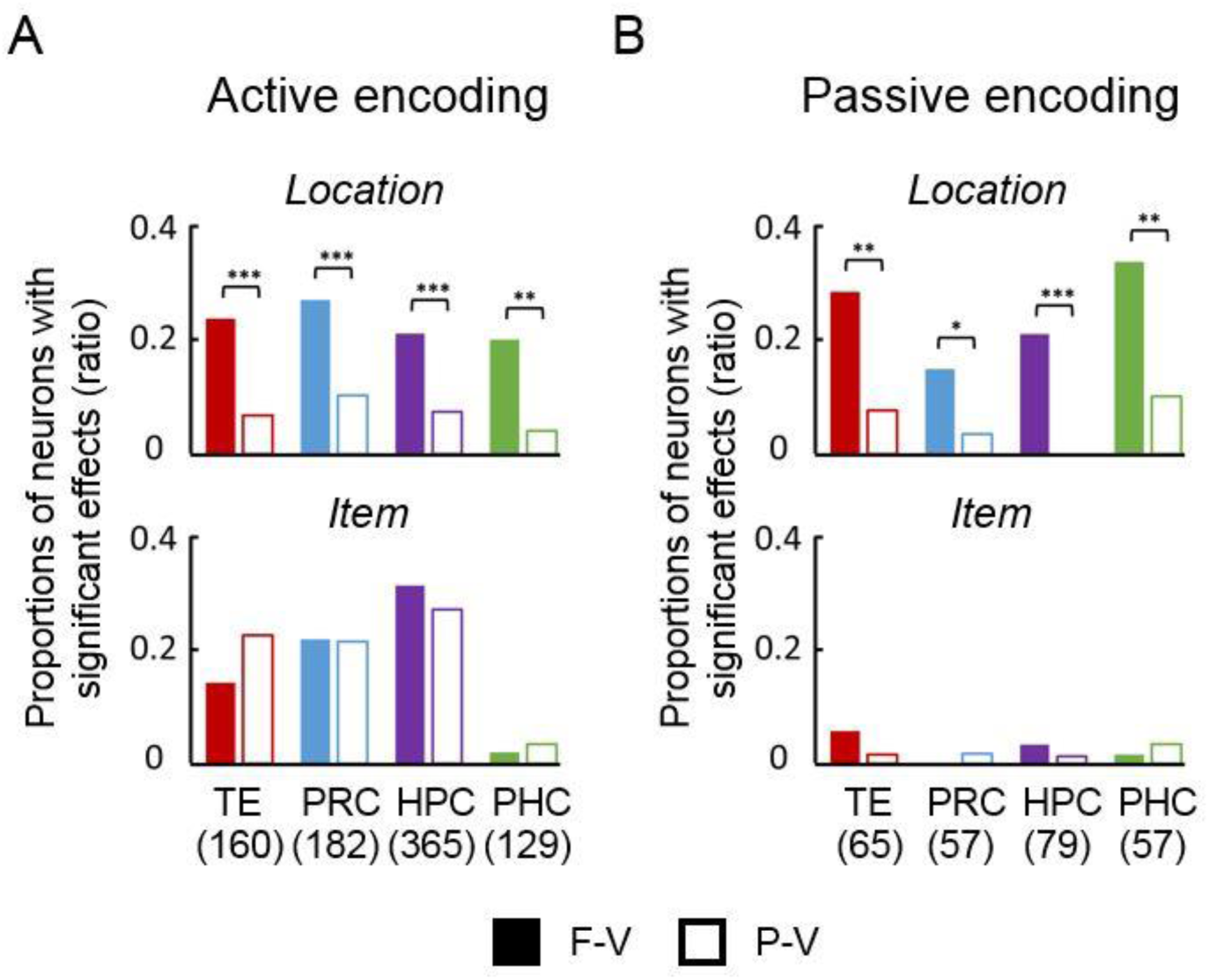
Proportions of location-selective and item-selective cells. (A) Proportions of location-selective cells (Top) and item-selective cells (Bottom) during the sample period (80-1000 ms after sample on) in the F-V (filled bars) and P-V conditions (open bars) in the active-encoding task. Numbers of recorded neurons (tested in both view conditions) are indicated in parentheses. **P < 0.0016, χ2 = 10.0 for PHC, d.f. = 1. ***P < 0.0001. χ2 = 19.5, 20.0, and 28.3 for TE, PRC, and HPC, respectively. (B) Proportions of location-selective cells (Top) and item-selective cells (Bottom) during the sample period in the F-V (filled bars) and P-V conditions (open bars) in the passive-encoding task. *P < 0.026, χ2 = 4.9 for PRC, d.f. = 1. **P < 0.005. χ2 = 11.1 and 8.7 for TE and PHC, respectively. ***P < 0.0001. χ2 = 20.3 for HPC.

The different sensitivity to the two view conditions was also observed for the location signal in the passive-encoding task (Figs. 2C&D). Similar to the active-encoding task, we found a substantial number of neurons exhibiting location effect (29%, TE; 15%, PRC; 21%, HPC; 34%, PHC) under the F-V condition (Fig. 3B). This result indicates that the location-selective response in the active-encoding task did not result from the task requirement, in which the animal was required to maintain actively a location of a sample stimulus. Compared with the F-V condition, the number of location-selective cells decreased dramatically under the P-V condition in all areas (8%, TE; 4%, PRC; 0%, HPC; 10%, PHC) (Fig. 3B). These results are also consistent with the single-unit results in the active-encoding task, and suggest that the gaze-sensitive location signal is automatically encoded by neurons in the TEv and MTL. The marked reduction of location signal in the P-V condition during either active or passive-encoding task argued against the possibility that the location-selective cells distinguish the structural organization of large objects with internal structures (e.g., a large grey square with a small letter at its top-left vs. at its bottom-right) which would be represented by the relational rather than the retinotopic space (Connor & Knierim, 2017).

The most straight-forward interpretation of fewer active location-selective cells under the P-V condition may be that fixating on the center of the display reduces attention to a sample stimulus and attenuates the response of location-selective cells, which showed robust location signals in the F-V condition. If this situation applies, we would then expect that neurons with stronger location selectivity in the F-V condition would show relatively stronger location selectivity in the P-V condition (i.e., a positive correlation). To test this possibility, we estimated strengths of location signals for neurons with location-selective activity in either F-V or P-V condition using *F* values indicating a location effect in the two-way ANOVA. Notably, we observed a negative correlation in amplitudes of the *F* values between the conditions in all areas during either active-encoding (Spearman rank correlation = −0.24 among 229 neurons across areas, *P* = 0.0003, two-tailed) (Fig. 4A) or passive-encoding task (Spearman rank correlation = −0.20 among 71 neurons, *P* = 0.090, two-tailed) (Fig. 4B). These results suggest that the weak location signal in the P-V condition was not due to the attenuated attention to a sample item. A reasonable interpretation of the negatively correlated location signal might be that separate visual inputs on the retinae drive different ensembles of neurons between the two view conditions (Fig. 1D). This interpretation is consistent with the significant reduction in the proportion of location-selective cells from the F-V to the P-V condition (Fig. 3) because a retinotopic shift of a large background square (48°, each side, Fig. 1C) in the F-V condition (Fig. 1D, left) would drive more neurons than that of a small sample stimulus (3°, radius) in the P-V condition (Fig. 1D, right). Collectively, the TEv and MTL areas may automatically signal large-scale background information represented on the retinotopic space, which necessarily reflects a perspective that a subject is viewing.

**Fig. 4.**
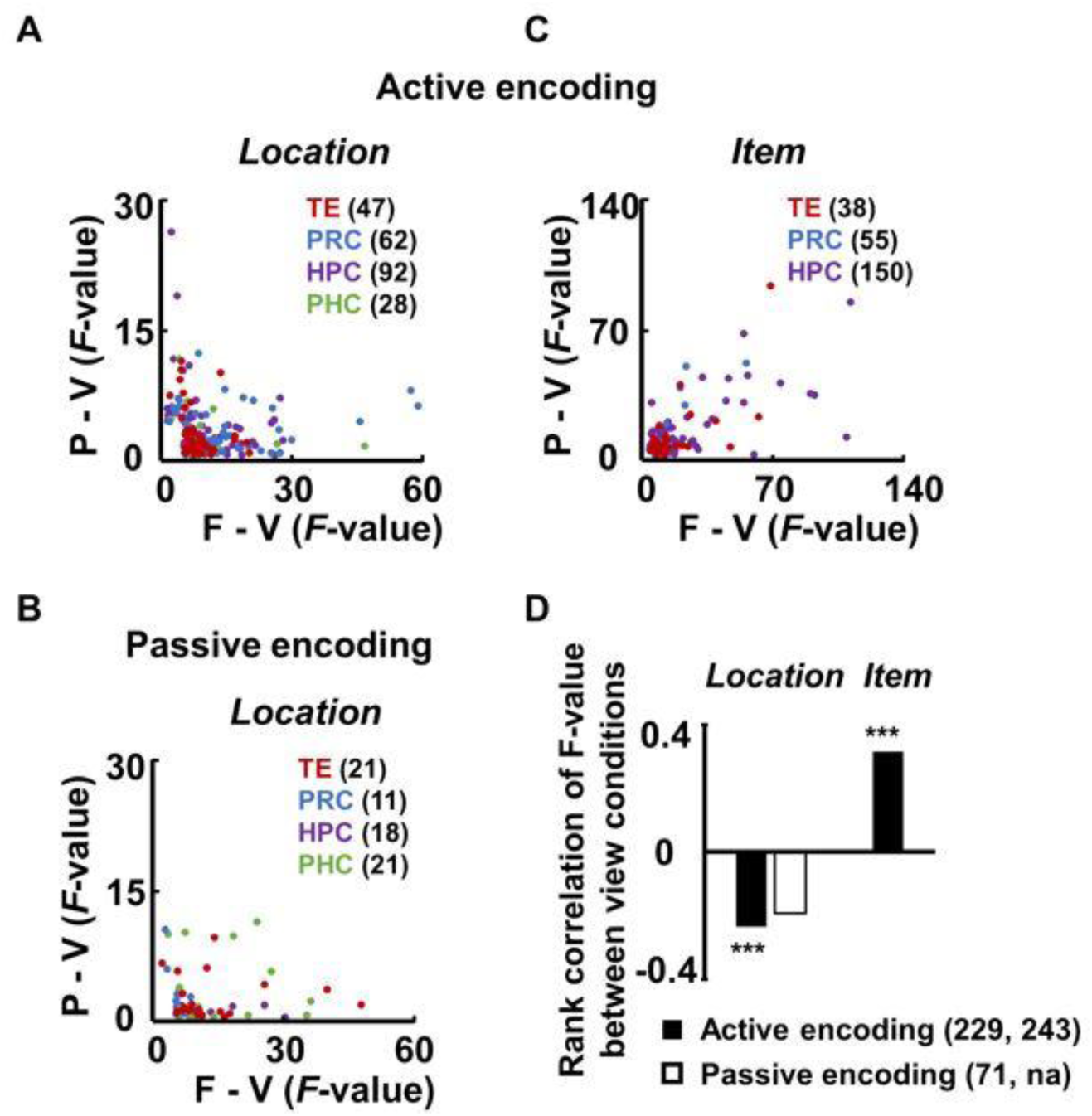
Location and item signal intensity between the two view conditions. (A) Location effect of the location-selective cells in the F-V and P-V conditions of the active-encoding task. F values in the P-V condition are plotted against those in the F-V condition for location-selective cells in either of the two view conditions. Neurons showing significant effects in either of the two conditions were used for the calculation of the F values. Numbers of the location-selective cells used for final calculation in each region are indicated in parentheses. (B) Location effect in the F-V and P-V conditions of the passive-encoding task. (C) Item effect of the item-selective cells in the two view conditions of the active-encoding task. The axis ranges in A-C were adjusted for display purpose, which included majorities of the data sets (A: 97.8%, B: 98.6%, C: 99.6%). (D) Correlation of the signal intensity between the two view conditions. Data from MTL and TEv were merged in the active-encoding and passive-encoding tasks, respectively. The total numbers of location-selective and item-selective cells used for final calculation are indicated in parentheses (left and right, respectively). na, not accountable. P = 0.0003, 0.0000(3.4E-07) and 0.09; ρ = −0.24, 0.32, and −0.20; d.f. = 227, 241 and 69 for the active-encoding (location), active-encoding (item), and passive-encoding (location), respectively. Spearman’s rank correlation, two-tailed.

### Task-dependent item signal

In contrast to the dramatic difference in the location-selective activity between the F-V and P-V conditions, neurons in the temporal lobe showed consistent item-selective responses between the two view conditions during the active-encoding task (Fig. S1). In all recording regions except for the PHC, we found a substantial number of item-selective cells under the P-V condition (TE 23%, PRC 22%, HPC 27%, and PHC 2%) as well as F-V condition (TE 14%, PRC 22%, HPC 32%, and PHC 3%) (Fig. 3A, bottom). These results are consistent with previous studies indicating the spatial invariance of object representation (Kobatake & Tanaka, 1994; Miyashita & Chang, 1988; Nakamura et al., 1994), which would be obtained by transforming it from the retinotopic space into the relational space along the ventral pathway (Connor & Knierim, 2017). In contrast to the location signal, the signal strengths of item information positively correlated between the F-V and P-V conditions (Figs. 4&D). These results indicate distinct processing between the item and its background (i.e., location signal) regarding their sensitivity to the view conditions. Interestingly, the number of item-selective cells was negligible in all areas under both view conditions in the passive-encoding task (F-V condition: TE 6%, PRC 0%, HPC 2%, PHC 3%; P-V condition: TE 2%, PRC 2%, HPC 3% PHC 1%; Fig. 3B, bottom), which contrasts to the substantial number of item-selective cells in the active-encoding task. The inconsistency in the item signal between the two tasks suggests that the object representation depends on the task demand, which required the subject to maintain an item identity of a sample stimulus for the following action.

### Population-coding analysis

The analyses based on the spike-firing data of individual neurons indicated substantially stronger location signal in the F-V condition compared with the P-V condition regardless of the task demands. One remaining question might be whether the location signal could be represented equivalently between the two view conditions by population coding. To test this possibility, we conducted the “representational similarity analyses” (RSA) (Kriegeskorte, Mur, & Bandettini, 2008); we first constructed a population vector consisting of firing rates of all recorded neurons in each area as its elements. In each combination of view condition and encoding-type, there were twenty-four (six items × four locations) of *n*-dimensional population vectors. “*n*” indicates a number of the recorded neurons in each area. We then calculated correlation coefficients between the population vectors, indicating the similarity level of neural representations between trial-types with different item-location combinations. Figures 5A and 5B displayed the similarity level of neural representations in the HPC during the sample presentation period in the active-encoding and passive-encoding tasks, respectively. In both tasks, the representational similarities between trial-types with same locations (e.g., location 1 item 1 & location 1 item 2) were substantially larger than the similarities between trial-types with different locations (e.g., location 1 item 1 & location 2 item 2) in the F-V condition (*P* <0.001 in both tasks, one-side, simulation test), suggesting that the HPC represents the item location that the animals were viewing, regardless of the task demands. In contrast to the F-V condition, the HPC’s discriminability in the location of a sample stimulus was considerably diminished in the P-V condition (Figs. 5A&B). In the RSA, other recorded regions also showed the marked reduction of the location signal in the P-V condition compared with the F-V condition in both tasks (Figs. 5C&D). Together, consistent with the analyses based on the single neurons, the analyses examining the population coding suggest that the temporal lobe areas represent the location information more robustly in the F-V condition than the P-V condition. As to the item signal, the RSA also provided the results which were consistent with the results of the single-neuron-based analyses (Fig. S2).

**Fig. 5.**
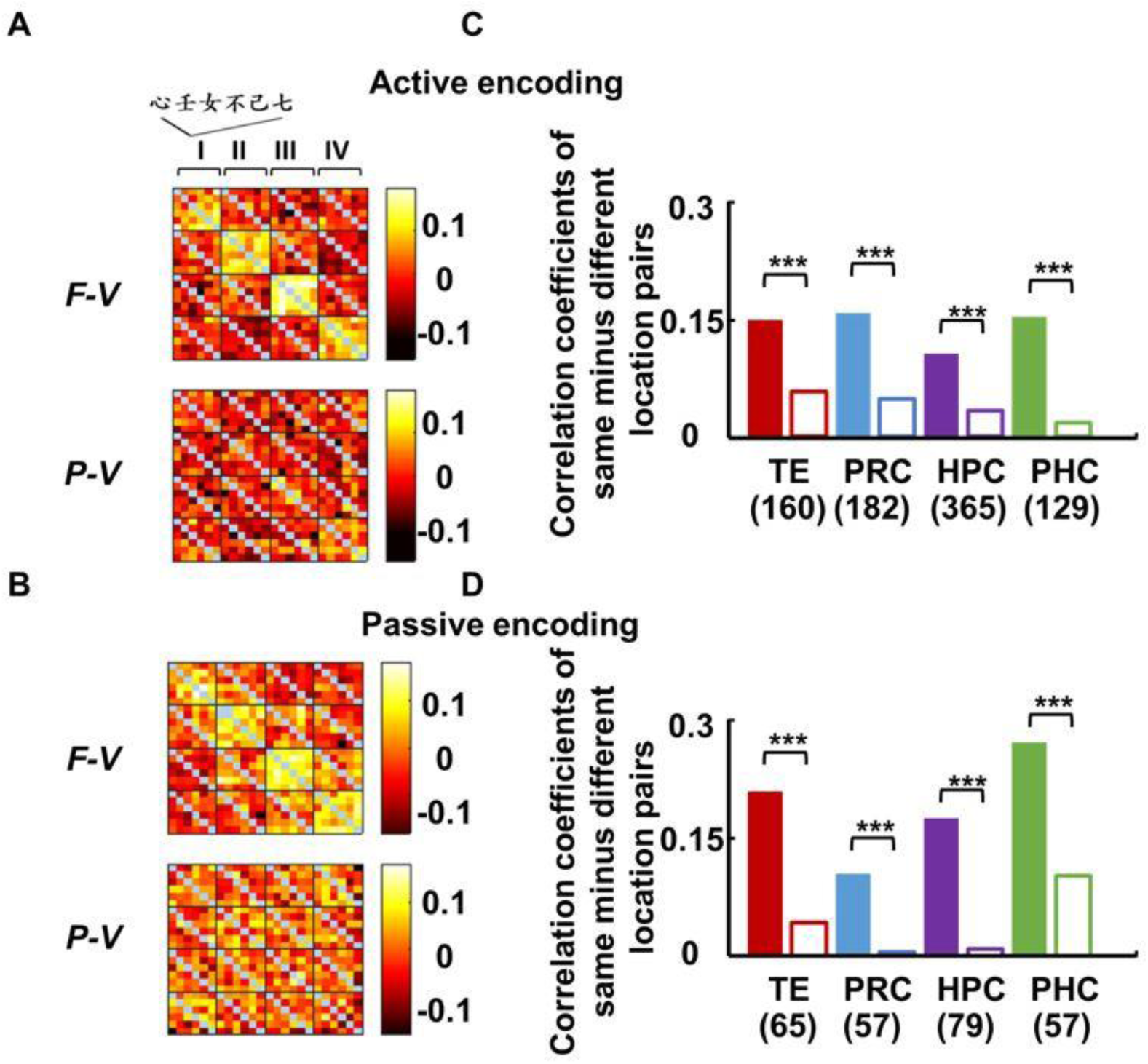
Location effects at population level. (A-B) Correlation coefficients of each pair out of the full 24 (four locations × six items)* 24 (four locations × six items) population vectors in the HPC under the F-V and P-V conditions of the (A) active-encoding and (B) passive-encoding tasks. Correlation coefficients of dummy data sets with location labels randomly shuffled (n=1000) were subtracted from the raw correlation coefficients. All recorded neurons from HPC were used in this analysis. Pearson’s linear correlation coefficient. (C-D) Difference value between the mean correlation coefficient under the same and different location pairs in the F-V and P-V conditions of the (C) active-encoding and (D) passive-encoding tasks in each brain region. The correlation coefficients between the population vectors for the trial-types with the same items (i.e., a diagonal line of each small matrix sorted by the locations, blue pixels in Figs. 5A&B) were excluded from this analysis.

### LFP activity depending on both view-condition and task-demand

In addition to spiking data, we investigated the LFP activity during the sample period. Figure 6A shows the differential spectrums between the viewing conditions (F-V condition minus P-V condition) in each recording region under the active-encoding task (*left column*) and passive-encoding task (*right column*). During the early sample presentation period (0-300 msec after sample onset), there is an enhanced beta-band activity (1-25 Hz) expressed non-selectively across the brain regions and tasks (Fig. 6B). This higher beta-band activity in the F-V condition is consistent with preceding literature indicating that larger beta-band activity is observed when the current cognitive or perceptual status should be actively maintained (i.e. the sample stimulus appears at the same position as with the fixation period in the F-V condition) than when the current state is disrupted by an unexpected event (i.e. the sample stimulus appears randomly at one out of the four positions in the P-V condition) (Engel & Fries, 2010). A view-condition dependent LFP activity was also observed in a gamma-band (30-80 Hz) during the late sample presentation period (350-800 msec after sample onset) (Fig. 6A). In contrast to the widely distributed beta-band, the gamma-band activity was selectively expressed only in the PRC and HPC when a sample item and its location were encoded actively by the foveal vision (Fig. 6B), in which situation both the item and location signals appeared robustly in these brain regions (Figs. 3&5&S2). These results may implicate that the increased gamma-band activity is related with the interaction between the item and location signals, which reportedly occurs in the PRC and HPC but not in TEv nor PHC (Chen & Naya, 2019).

**Fig. 6.**
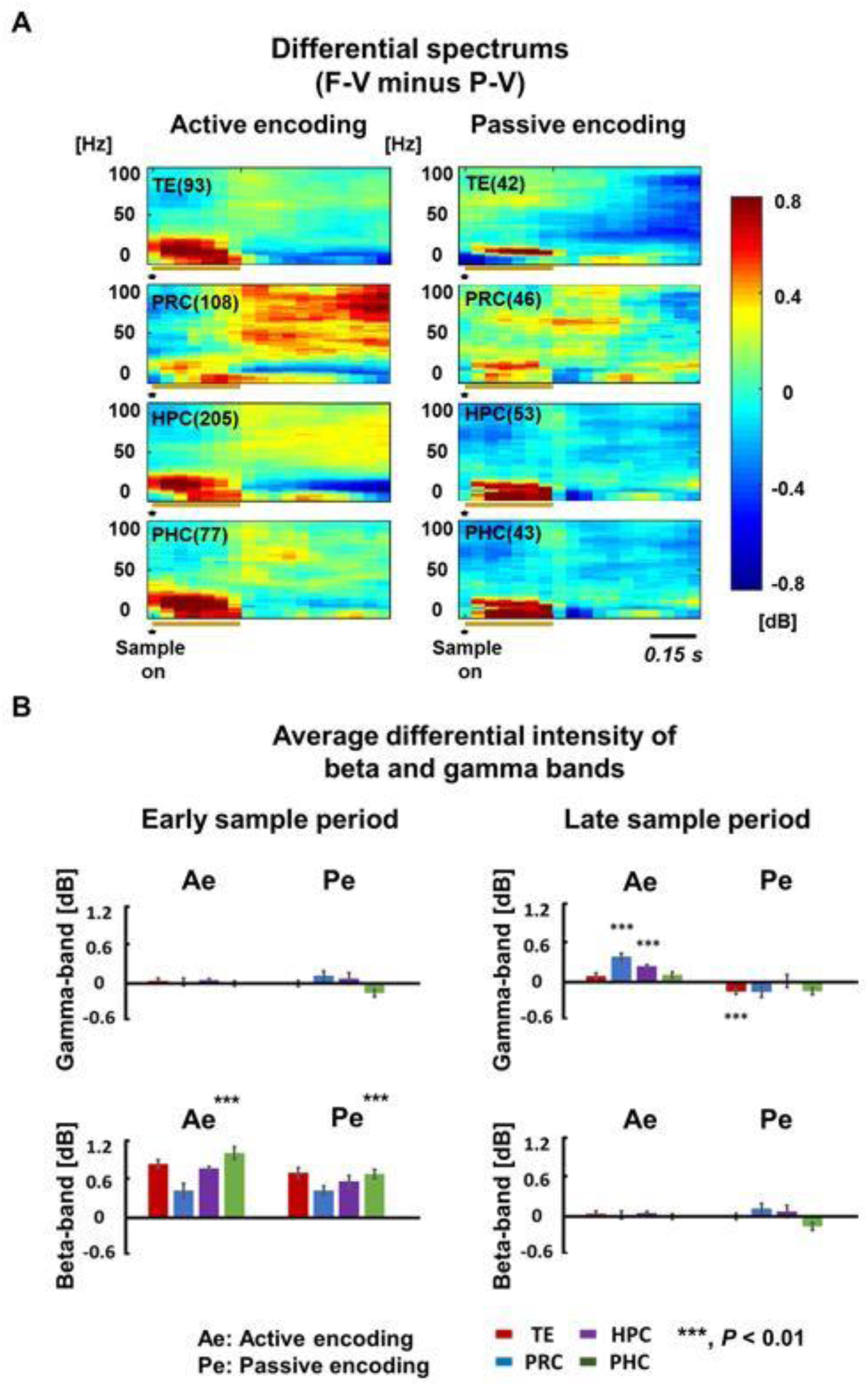
The difference spectrums of the local field potential activities between the F-V and P-V conditions in the active-encoding and passive-encoding tasks. (A) The average difference local field potential (LFP) spectrums between the F-V and P-V conditions of active-encoding and passive-encoding tasks during the sample period (0:1000 msec after sample on). Raw spectrums from different recording sites (indicated in parentheses) were used for this analysis. Average intensity of each frequency during the baseline period (600:0 msec before sample on) was subtracted at the corresponding frequency. (B) The average difference value of beta-band (1-25 Hz) and gamma-band (30-80 Hz) intensity during early sample (0:300 msec after sample on) and late sample (350:800 msec after sample on) periods (see Fig. 6A) in each task and recording region.

## Discussion

The present study provides single-unit data showing robust spatial information in the TEv and MTL areas, which signaled a particular location where the animals were viewing (F-V condition) rather than an object position presented in the peripheral view (P-V condition). These results were shown for each of the recording regions by the independent analyses for each of the two monkeys, indicating the very robust animal consistency. In addition, this animal consistency was confirmed even though the two animals were tested in different task demands (i.e., active-encoding and passive-encoding of an object and its location), which manifests the robustness of the present findings showing an existence of the location signal characterized by the clear difference in its sensitivity to t he two view conditions. These new findings suggest that the location signal in the primate temporal lobe areas may represent a view-centered background image, which could specify the current gaze position within a scene (Fig. 7). This view-centered background may be automatically represented in the temporal lobe areas because it was observed in the passive-encoding task as well as the active-encoding task. The TEv and MTL areas except for the PHC also signaled object information. However, in contrast to the background information, the object information was represented regardless of the view conditions when it was actively encoded. These results from the single-neuron-based analyses were confirmed by population-coding analyses. Taken together, the present study suggests that the ventral pathway and its downstream in the MTL signal not only spatially-invariant object information but also view-centered background information, which may automatically locate the object in a scene when it is viewed by the foveal vision.

**Fig. 7.**
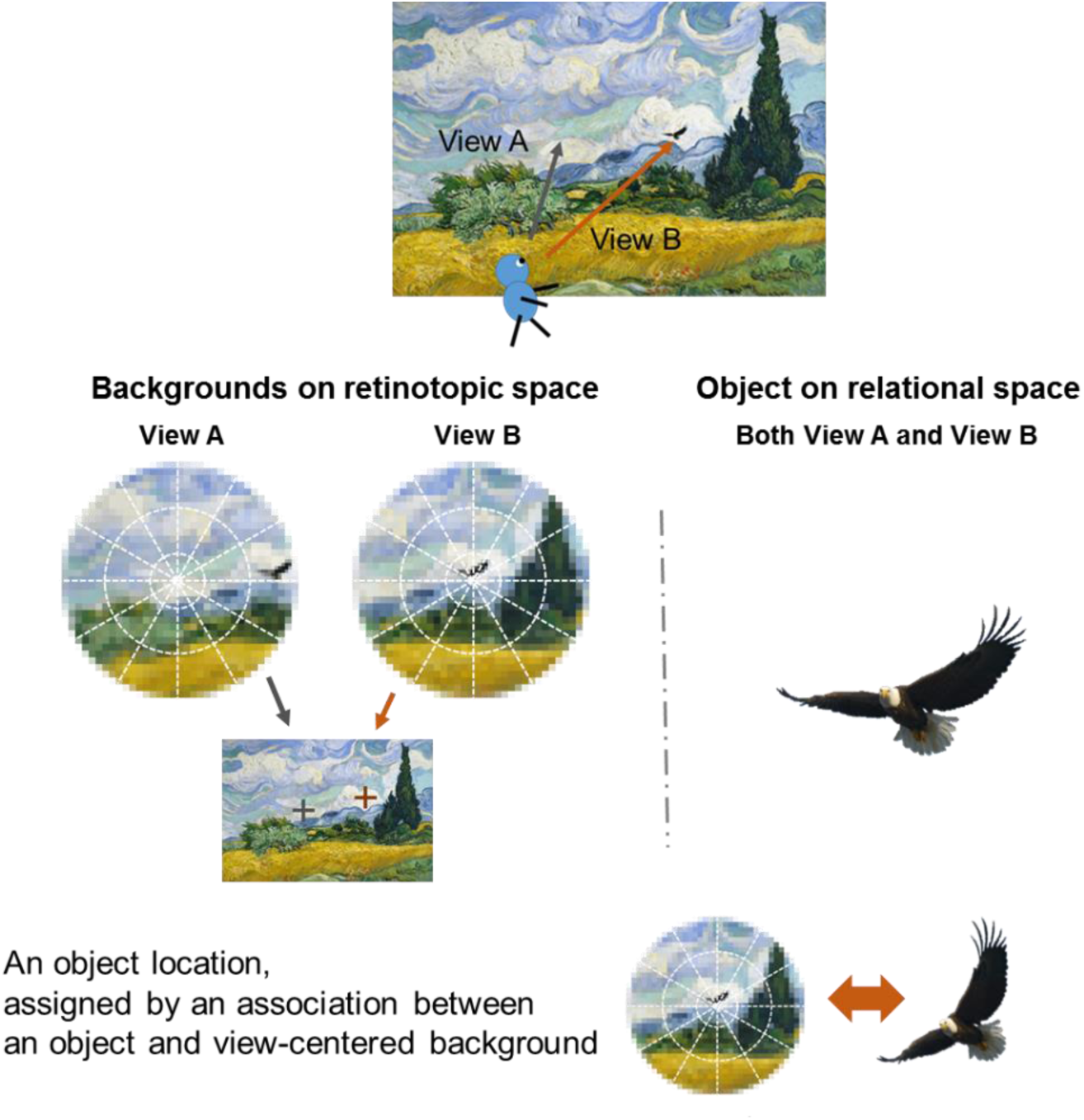
Parallel scene processing on the retinotopic and relational spaces. *Top,* Assume that a subject was in wheat field and viewing the valley. An eagle was in parafoveal vision of the subject in view A, while it was in the subject’s foveal vison in View B. *Middle*, When the subject attended the eagle either voluntarily or involuntarily, the eagle would be selected as an object from the retinotopic image and processed on the relational space (*right*) regardless of its original retinotopic. Conversely, background images would be automatically captured and processed on the retinotopic space, which specify a location of the view point in the scene accordingly (*left*). *Bottom*, The location of the object in the scene would be assigned by an associated information between the view-centered background and the object. This model hypothesizes that first person’s perspective of a scene containing objects depends on the parallel visual processing on the retinotopic and relational spaces, and their association. The original painting is titled Wheat Field with Cypresses by Vincent Willem van Gogh.

One naïve question on the gaze-related location signal might be whether the location signal could be explained by non-visual sensory/motor information, which reflects the animals’ eye positions relative to their heads. Our previous study indicated that neurons in the TEv and MTL areas responded differently to the same gaze positions depending on the position of the large background square within the display (leftward or rightward) (Chen & Naya, 2019), suggesting that the gaze-related location signal reflects visual inputs rather than somatosensory/motor-related information of the gaze itself. In the present study, we characterized the location signal, which were widely distributed over the temporal lobe areas, by revealing the underlying visual inputs not to be represented on the relational space, but instead on the retinotopic space (i.e., view-centered background). An important question about the view-centered background information on the retinotopic space might be whether it only reflects the parafoveal vision or not. In the present study, the location-selective activity depends on the parafoveal vision of the background, which shows an edge of the large grey square or the display frame. However, some neurons exhibited location-selective activities only after sample presentation in the F-V condition (Figs. 2B-D) (10.8% and 8.5% across areas in the active and passive-encoding tasks), which suggest an existence of neuronal population that represent the view-centered background including foveal vision as well as parafoveal vision. The view-centered background signal in the present study may explain response patterns of “spatial view cells” in the HPC (and posterior PRC) reported by Rolls (Rolls, Robertson, & Georges-François, 1997). The spatial view cells show selective responses to a particular location where an animal views regardless of its standing position. This allocentric coding property of the spatial view cells could be due to similar visual inputs when an animal views the same location from different positions.

In spite of the location signal which may reflect the background information on the retinotopic space, the object signal was detected regardless of its retinotopic position in the active encoding task (Fig. 7), which confirmed the preceding literature showing the spatial invariant of object representation in the IT cortex (Miyashita & Chang, 1988; Nakamura et al., 1994). The representation of an object may be explained by a spatial relationship among the internal elements of it, which necessarily accompany its transformation from the retinotopic space into the relational space (Connor & Knierim, 2017). The present study suggests that neurons in the temporal lobe signal the location information of an object as its background image represented on the retinotopic space (Fig. 7) rather than an interrelation between the object and any other spatial structure such as a large gray square behind it. Based on the present experimental set up, the background image encoded by neurons in the TEv and MTL areas should cover larger than 30 degrees in the visual angle (diameter) to include the edge of the large gray square background, which may cause different responses according to the gaze positions. As well as the object signal, the large-scale background image is reportedly processed along the ventral pathway (Kornblith et al., 2013; Vaziri et al., 2014). One remaining question is whether the processing of the background image in the ventral pathway imparts more generalized spatial features (e.g., field, valley, forest), which may be represented on the relational space and serve for recognizing an entire scene (e.g., suburb rather than modern city) regardless of the gaze positions (e.g., an eagle over the valley).

In addition to the view conditions testing the representation spaces (i.e., relational vs. retinotopic), the object and the background signals showed differential sensitivity patterns to the task demands in the present study. The background signal was encoded irrespective of the task demand while the object signal was encoded only in the active-encoding task. The automatic encoding of the background signal suggests that when we direct our gaze toward an object to obtain its high-resolution image, we would spontaneously receive the spatial information, which would be assigned to the object (Chen & Naya, 2019). One remaining problem about the object signal might be whether the lack of item-selective activity in the passive-encoding task is due to the present stimulus set (i.e., Chinese character) because the IT neurons reportedly respond to object stimuli such as face stimuli in a passive-viewing task (Kiani, Esteky, Mirpour, & Tanaka, 2007; Tsao, Freiwald, Knutsen, Mandeville, & Tootell, 2003). Compared with a natural object such as a face stimulus, a fabricated two-dimensional stimuli used in the present study may not bring about a bottom-up attention to be perceived as an object. In the active-encoding task, the monkey learned the Chinese characters to discriminate one from another. The repetitive training in the active-encoding task might form a long-term learning effect on the stimulus to induce the bottom-up attention, which may lead a transformation of representations of Chinese-characters from the retinotopic space into the relational space. Although we cannot address if the attention was derived from the bottom-up or the top-down, the attention-dependent object signal and the attention-independent background signal may derive from a figure-background segmentation, which reportedly occurred at the V4, a start point of the ventral pathway (Roe et al., 2012). Previous studies have focused on the object information which is filtered, and implicated that the object representation is transformed from the retinotopic space into the relational space with the increase of neurons’ receptive fields along the ventral pathway (Connor & Knierim, 2017). We hypothesize that the background information, which is filtered-out at the figure-ground segmentation, spreads into the ventral pathway with its representation remaining on the retinotopic space rather than the relational space. Our previous report has demonstrated that the two distinct signals, which are segmented from the same retinal image, are integrated step-by-step from the TEv, PRC to HPC (Chen & Naya, 2019). From the ventral stream to the MTL areas, the strongest integration effect was found in the PRC at the single neurons level. This integration process may be related with the largest gamma-band LFP activity in the PRC, which was observed when the monkey gazed at an object to encode its identity and location information actively (Fig. 6).

In the present study, the PHC represents the view-centered background signal whose property is similar to that in the TEv and other MTL areas including the PRC. Considering the heavier projections from the posterior parietal cortex to the PHC compared with the AIT cortex including the PRC (Kravitz, Saleem, Baker, & Mishkin, 2011), the PHC may also process the spatial information related with the eye/self-movement. Contributions of the PHC to scene construction process may become apparent when a subject perceives the environment by moving their gazes (Zhang & Naya, 2019) in which multiple views should be coordinated according to the eye/self-movements, beyond encoding a single snapshot focusing on one object which was investigated in the present study. We propose a future study to investigate how the past multiple views influence on the present view to build the current first person’s perspective (Eichenbaum et al., 2007; Palombo et al., 2015; Tulving, 2002), which may be related with an encoding of episodic memory.

## Materials and Methods

### Subjects

Two male monkeys (*Macaca mulatta*) (9.3 kg, monkey A; 10.1 kg, monkey F) were used for the experiments. All procedures and treatments were performed in accordance with the NIH Guide for the Care and Use of Laboratory Animals and were approved by the Institutional Animal Care and Use Committee (IACUC) of Peking University.

### Behavioral task

We trained monkey A on a foveal-view/F-V condition of an active-encoding task with six visual items (Fig. 1). During both training and recording sessions, monkeys performed the task under dim light in an electromagnetic shielded room (length * width * height = 160 cm *120 cm * 222 cm). The task began with an encoding phase, which was initiated by the animal pulling a lever and fixating on a white square (0.6 ° of visual angle) presented within one of the four quadrants (12.5 ° from the center) of a touch screen (3MTM MicroTouchTM Display M1700SS, 17 inch, horizontal viewing angle: ∼59 °, vertical viewing angle: ∼49 °) with a custom-made metal frame (diagonal size: 22 inch, horizontal viewing angle: ∼72 °, vertical viewing angle: ∼71 °) situated ∼28 cm from the subjects. Eye position was monitored using an infrared digital camera with a 120 Hz sampling frequency (ETL-200, ISCAN) placed next to the left edge of the touch screen. The eye position calibration was conducted before starting each recording session (Monkey logic). After a 0.6 s fixation, one of the six items (3.0 °, radius) was presented in the same quadrant as a sample stimulus for 0.3 s, followed by another 0.7 s fixation on the white square. An additional 0.017 s, reflecting the design of software and hardware controlling the behavioral task was added to each trial event. If the fixation was successfully maintained (typically, < 2.5 °), the encoding phase ended with the presentation of a single drop of water.

The encoding phase was followed by a blank interphase delay interval of 0.7-1.4 s during which no fixation was required. Then, the response phase was initiated with a fixation dot presented at the center of the screen. One of the six items was then presented at the center for 0.3 s as a cue stimulus. After another 0.5 s delay period, five discs were presented as choices, including a blue disc in each quadrant and a green disc at the center. When the cue stimulus was the same as the sample stimulus, the subject was required to choose by touching the blue disc in the same quadrant as the sample (i.e., match condition). Otherwise, the subject was required to choose the green disc (i.e., nonmatch condition). If the animal made the correct choice, four to eight drops of water were given as a reward; otherwise, an additional 4 s was added to the standard intertrial interval (1.5-3 s). During the trial, a large gray square (48 ° on each side, RGB value: 50, 50, 50, luminance: 3.36 cd/m^2^) was presented at the center of the display (backlight luminance: 0.22 cd/m^2^) as a background. After the end of a trial, all stimuli disappeared and the entire screen displayed light-red color during the inter-trial interval. The start of a new trial was indicated by the re-appearance of the large gray square on the display, upon which the monkey could start to pull the lever triggering an appearance of a white fixation dot. In the match condition, sample stimuli were pseudorandomly chosen from six well-learned visual items, and each item was presented pseudorandomly within the four quadrants, resulting in 24 (6 × 4) different configuration patterns. In the nonmatch condition, the position of the sample stimulus was randomly chosen from the four quadrants, and the cue stimulus was randomly chosen from the five items that differed from the sample stimulus. The match and nonmatch conditions were randomly presented at a ratio of 4:1, resulting in 30 (24+6) different configuration patterns. The same six stimuli were used during all recording sessions.

In addition to the F-V condition, we tested the neuronal responses of monkey A in the peripheral-view/P-V condition of the active-encoding task. In this view condition, fixation on the center of the display was required during the encoding phase (Fig. 1). Other parameters were the same as those in the F-V condition of the active-encoding task. Correct performance under F-V condition: 97.5 ± 2.6% in the match trials and 90.8 ± 8.1% in the nonmatch trials (n = 454 sessions); P-V condition: 94.3 ± 6.2% in the match trials and 84.1 ± 10.8% in the nonmatch trials (n = 478 sessions).

We tested the neuronal responses of monkey F in both F-V and P-V condition of a passive-encoding task, in which the task sequence and requirement were same as the encoding phase of the active-encoding task but without a lever-pulling requirement (no interphase delay interval and response phase). The configuration of visual stimuli (such as visual angles, configuration patterns, and others) was same as that for monkey A. We tested the neuronal response of both monkey A and monkey F in the F-V and P-V conditions in a block manner.

### Electrophysiological recording

Following initial behavioral training, animals were implanted with a head post and recording chamber under aseptic conditions using isoflurane anesthesia. To record single-unit activity, we used a 16-channel vector array micrILRobe (V1 X 16-Edge, NeuroNexus), 16-channel U-Probe (Plexon), tungsten tetrode probe (Thomas RECORDING), or a single-wire tungsten microelectrode (Alpha Omega), which was advanced into the brain using a hydraulic Microdrive (MO-97A, Narishige) (Naya & Suzuki, 2011). The microelectrode was inserted through a stainless steel guide tube positioned in a customized grid system on the recording chamber. Neural signals for single units were collected (low-pass, 6 kHz; high-pass, 200 Hz) and digitized (40 kHz) (OmniPlex Neural Data Acquisition System, Plexon). These signals were then sorted using an offline sorter provided by the OmniPlex system. We did not attempt to prescreen isolated neurons. Instead, once we isolated any neuron, we started to record its activity. The location of microelectrodes in target areas was guided by individual brain atlases from MRI scans (3T, Siemens). We also constructed individual brain atlases based on the electrophysiological properties around the tip of the electrode (e.g., gray matter, white matter, sulcus, lateral ventricle, and bottom of the brain). The recording sites were estimated by combining the individual MRI atlases and physiological atlases (Naya, Chen, Yang, & Suzuki, 2017). To record LFPs, we used neural signals from the same electrodes as we used for the recording of spikes. However, the signals were collected using different filters (low-pass, 200 Hz; high-pass, 0.05 Hz), and digitized at 1 kHz.

The recording sites in monkey A covered an area between 5 and 24 mm anterior to the interaural line (right hemisphere). The recording sites in monkey F covered an area between 6.6 and 23.4 mm anterior to the interaural line (right hemisphere). The recording sites in HPC appeared to cover all its subdivisions (i.e., dentate gyrus, CA3, CA1, and subicular complex). The recording sites in PHC focused on approximately the lateral 2/3. The recording sites in PRC appeared to cover areas 35 and 36 from the fundus of the rhinal sulcus to the medial lip of the anterior middle temporal sulcus (amts). The border of PRC’s caudal limit (PHC’s rostral limit) was determined according to the rostral limit of the occipital temporal sulcus and the caudal limit of the rhinal sulcus (Suzuki & Amaral, 2003). In monkey A, the caudal limit of the recording sites in PRC is 2 mm posterior to the caudal limit of its rhinal sulcus and 1 mm anterior to the rostral limit of the occipital temporal sulcus. In monkey F, the caudal limit of the recording sites in PRC is 0 mm posterior to the caudal limit of its rhinal sulcus and 0 mm anterior to the rostral limit of the occipital temporal sulcus. The recording sites in TE were limited to its ventral area, including both banks of the amts.

### Data analysis

All neuronal data were analyzed using MATLAB (MathWorks) with custom written programs, including the statistics toolbox. For responses before sample presentation, we tested each neuron’s firing rate during the 700 ms period before the sample stimulus onset, including the 100 ms before the fixation start, as the monkeys typically started fixation 160-170 ms after fixation dot presentation. For responses during/after sample presentation, the firing rate during the period extending from 80 to 1000 ms after sample onset was tested. For responses before sample presentation, we evaluated the effects of “location” for each neuron using one-way ANOVA (*P* < 0.01). For sample responses, we evaluated the effects of “location” and “item” for each neuron using two-way ANOVA with interactions (P < 0.01 for each). We analyzed neurons that we tested in at least 60 trials (10 trials for each stimulus, 15 trials for each location).

## Acknowledgments

We thank E.T. Rolls, W.A. Suzuki, M. Zhang, K.W. Koyano, C. Yang for helpful comments and S. Xue for expert animal care. Funding: The present study was funded by National Natural Science Foundation of China Grant 31421003 & 31871139 (to Y.N.).

## Author Contributions

Y.N. designed the experiments. H.C. performed the experiments. H.C. and Y.N. analyzed data and wrote the manuscript.

## Supplementary Figures

**Fig. S1.**
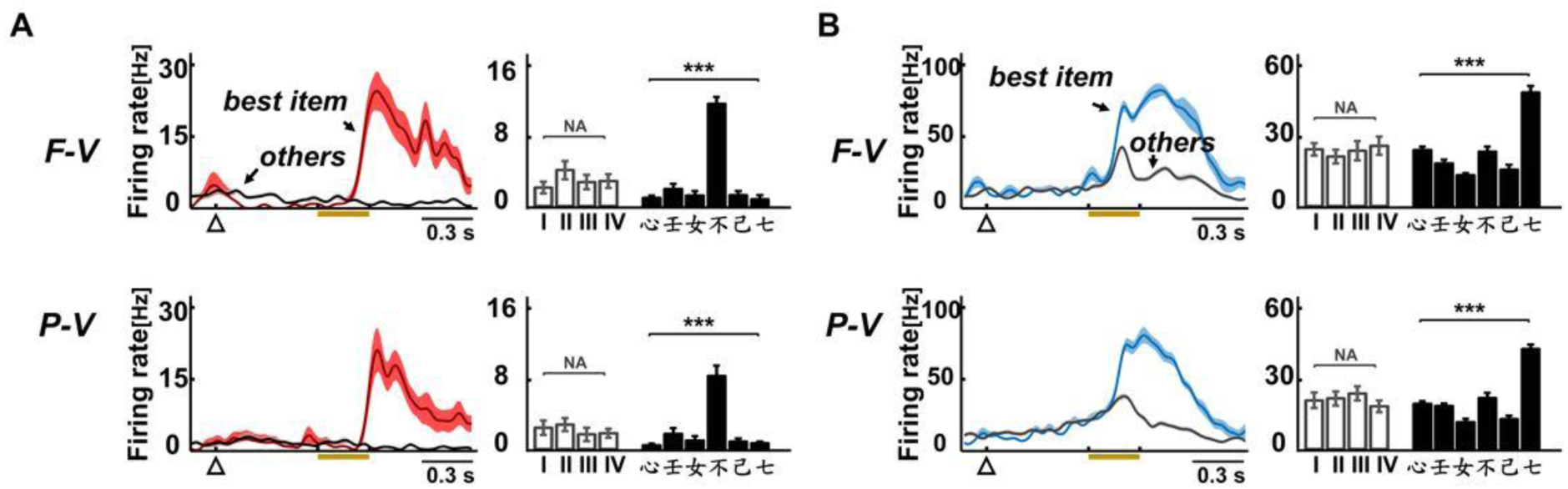
Responses of the item-selective cells in the active-encoding task. (A) Example of the item-selective cells from TE in the F-V and P-V conditions of the active-encoding task. (Left) Spike-density functions (SDFs) (sigma = 20 ms) indicating the firing rates under two conditions (best item and the average of other five items). (Right) Bar graph indicating the mean firing rate during sample period (80-1000 ms after sample on) under each location and each item. (B) Example of the item-selective cells from PRC in the F-V and P-V conditions of the active-encoding task.

**Fig. S2.**
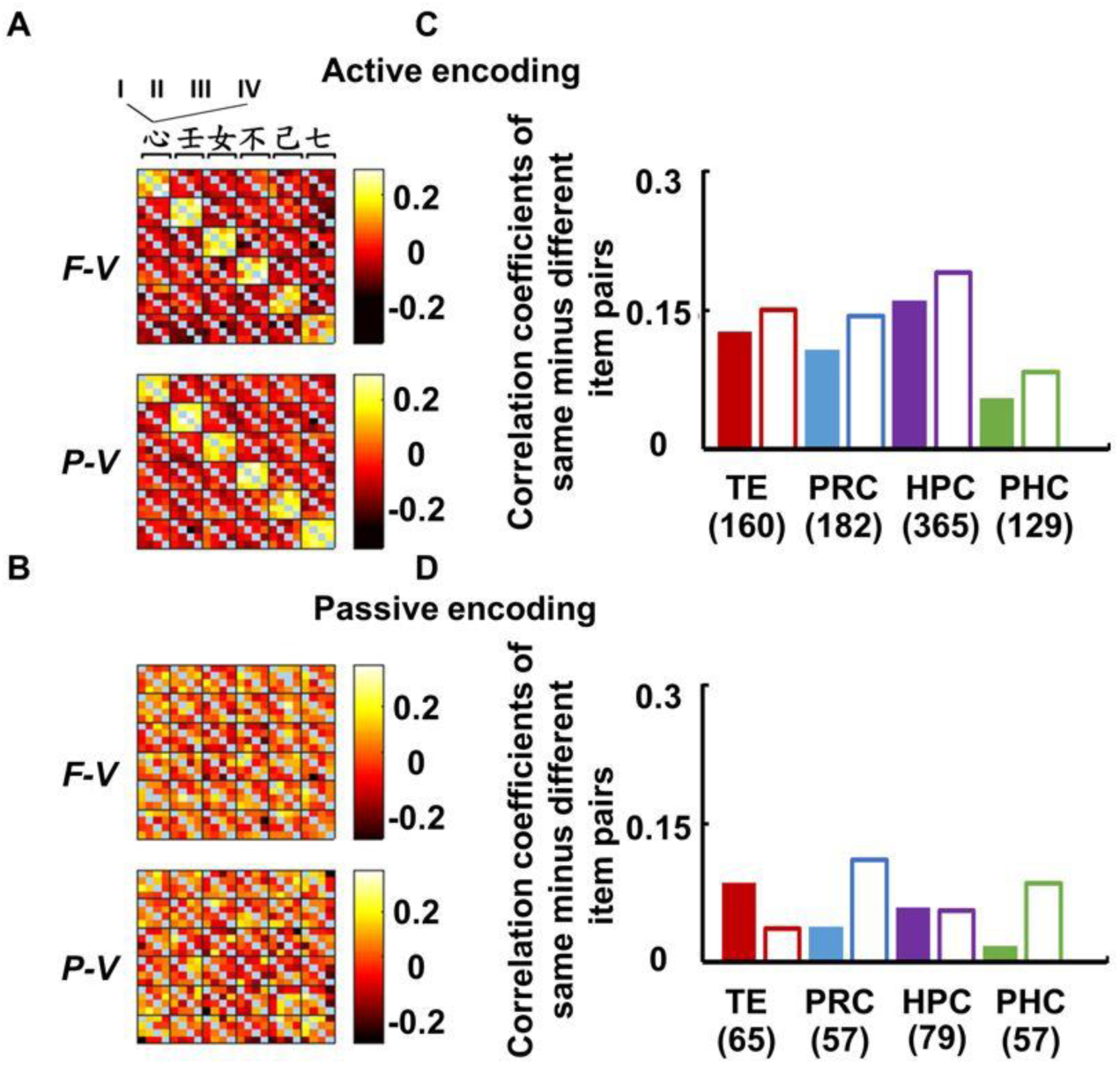
Item effects at population level. (A-B) Correlation coefficients of each pair out of the full 24 (six items × four locations) * 24 (six items × four locations) population vectors in the HPC under the F-V and P-V conditions of the (A) active-encoding and (B) passive-encoding tasks. Correlation coefficients of dummy data sets with item labels randomly shuffled (n=1000) were subtracted from the raw correlation coefficients. All recorded neurons from HPC were used in this analysis. Pearson’s linear correlation coefficient. (C-D) Difference value between the mean correlation coefficient under the same and different item pairs in the F-V and P-V conditions of the (C) active-encoding and (D) passive-encoding tasks in each brain region. The correlation coefficients between the population vectors for the trial-types with the same items (i.e., a diagonal line of each small matrix sorted by the items, blue pixels in Figs. S2A&B) were excluded from this analysis.

